# Impact of fluoroquinolones and aminoglycosides on *P. aeruginosa* virulence factor production and cytotoxicity

**DOI:** 10.1101/2021.10.11.463927

**Authors:** Daniel M. Foulkes, Keri McLean, Marta Sloniecka, Dominic Byrne, Atikah S. Haneef, Craig Winstanley, Neil Berry, David G. Fernig, Stephen B. Kaye

## Abstract

Infection from the opportunistic pathogen *Pseudomonas aeruginosa* is one of leading causes of disability and mortality worldwide and the world health organisation has listed it with the highest priority for the need of new antimicrobial therapies. *P. aeruginosa* strains responsible for the poorest clinical outcomes express either ExoS or ExoU, which are injected into target host cells via the type III secretion system (T3SS). ExoS is a bifunctional cytotoxin that promotes intracellular survival of invasive *P. aeruginosa* by preventing targeting of the bacteria to acidified intracellular compartments and lysosomal degradation. ExoU is a potent phospholipase which causes rapid destruction of host cell plasma membranes, leading to acute tissue damage and bacterial dissemination. Fluoroquinolones are usually employed as a first line of therapy as they have been shown to be more active against *P. aeruginosa in vitro* than other antimicrobial classes. However, their overuse over the past decade has caused alarming rates of antibiotic resistance to emerge. In certain clinical situations, aminoglycosides have been shown to be more effective then fluoroquinolones, despite their reduced potency towards *P. aeruginosa in vitro*. In this study, we evaluated the effects of fluoroquinolones (moxifloxacin and ciprofloxacin) and aminoglycosides (tobramycin and gentamycin) on T3SS expression and toxicity, in corneal epithelial cell infection models. We discovered tobramycin disrupted T3SS expression and inhibited both ExoS and ExoU mediated cytotoxicity, protecting infected HCE-T cells even at concentrations below the minimal inhibitory concentrations (MIC). Fluoroquinolones moxifloxacin and ciprofloxacin, however, upregulated the T3SS and in particular did not subvert the cytotoxic effects of ExoS and ExoU.

## Introduction

*Pseudomonas aeruginosa* is Gram-negative bacterium that colonises a diverse range of environmental niches. *P. aeruginosa* is also a major opportunistic pathogen and common cause of nosocomial infection, associated with a wide range of diseases, including pneumonia and microbial keratitis [1-3]. It is a leading cause of intensive care unit-acquired pneumonia (ICUAP) [4], and is the second most frequent colonising bacteria in patients with COVID19 [5, 6]. It is also the primary causative agent of bacterial keratitis, which is recognised as the second largest cause of legal blindness worldwide [7]. As a pathogen of current major concern, the world health organisation (WHO) has listed carbapenem-resistant *P. aeruginosa* (CRPA) with the highest priority for the development of new antimicrobial therapies [8].

Pathogenic *P. aeruginosa* strains use the type III secretion system (T3SS), to inject exotoxins directly into the cytoplasm of compromised host epithelia [9]. The T3SS has been identified as a principal virulence determinant for poor clinical outcomes in pneumonia, sepsis, keratitis, and otitis externa [2-4, 10, 11]. T3SS expressing *P. aeruginosa* clinical isolates can be further categorised as either exotoxin S (ExoS) or exotoxin U (ExoU) producing. In a study of hospitalised patients with *P. aeruginosa* bacteraemia, 97.5% of bloodstream isolates were positive for *exoU* or *exoS* genes, with isolates containing *exoU* being significantly more resistant to antibiotic treatment [12]. ExoS is a bifunctional cytotoxin, possessing dual Rho GTPase-activating protein (GAP) and ADP-ribosyltransferase activity and is a critical virulence factor enabling *P. aeruginosa* strains with invasive phenotypes to avoid being targeted to acidified intracellular compartments such as the lysosyme [13][14]. ExoU is a ubiquitin activated phospholipase that is localised to the inner leaflet of the host cell plasma membrane via phosphatidylinositol 4,5-bisphosphate (PIP_2_) dependent targeting, where it exhibits cytotoxicity by cleaving phospholipids, resulting in host cell lysis [15]. ExoU catalytic activity is directed towards phospholipids at the sn-2 position, and results in arachidonic acid release which induces pathways that result in NF-κB activation and MAPK signalling [16-18]. This leads to upregulation of IL-8 and keratinocyte chemoattractant (KC), and increased infiltration of neutrophils that exacerbate tissue damage via acute localised inflammation [16, 17].

T3SS expression and production of ExoS and ExoU is tightly controlled at the transcriptional level in response to environmental cues, including contact with host cells and low levels of extracellular calcium ions [18, 19]. Expression is controlled principally by the interactions of four transcription factors: ExsA, ExsC, ExsD, and ExsE, with the AraC family transcription factor, ExsA serving as the primary activator of *P. aeruginosa* T3SS gene expression [20]. ExsA DNA biding induces expression of several proteins that form the T3SS macromolecular complex, spanning the inner bacterial membrane, the periplasmic space, the peptidoglycan layer, the outer bacterial membrane, the extracellular space, and the host cell membrane [9]. The needle like structure is assembled by helical polymerisation of PscF proteins [21]. PcrV is an essential translocator protein which forms the needle tip, without which exotoxins cannot be secreted [22].

T3SS expression in *P. aeruginosa* is associated with acute toxicity, and delay of- or failure to initiate adequate antimicrobial therapy is linked to increased mortality [10]. Fluoroquinolones, such as moxifloxacin and ciprofloxacin, disrupt bacterial DNA replication by inhibiting DNA topoisomerases and DNA-gyrases, and are normally the primary line of treatment for *P. aeruginosa* infections [23]. They demonstrate high potency against most clinical isolate strains of *P. aeruginosa in vitro*, however, their use has led to the emergence of *P. aeruginosa* strains with resistant phenotypes, predominantly via efflux dependent mechanisms [24]. Aminoglycosides are another group of antimicrobial agent used in the treatment of *P. aeruginosa* infections, which function by binding to the A-site (aminoacyl) of 16S rRNA, a component of the bacterial ribosomal 30S subunit, to disrupt protein synthesis [25]. In comparison to fluoroquinolones, aminoglycosides (such as tobramycin, amikacin, and gentamicin) are generally considered less potent compounds towards *P. aeruginosa* when assayed *in vitro*; however, they often demonstrate improved utility against *P. aeruginosa* infections in certain clinical settings. For example, inhalation of aerosol formulations of aminoglycosides, especially tobramycin, have proven efficacious in the treatment and prevention of bronchiectasis [26]. Administration of tobramycin has also recently been shown to reduce the formation of neutrophil extracellular traps (NETs) in a murine model of *P. aeruginosa* lung infection [27]. Despite being a major determinant in disease progression and clinical outcome, the effects these antimicrobials have on T3SS expression (if any) is currently unknown. In this study we explore the mechanisms through which such antimicrobials might influence *P. aeruginosa* T3SS-dependent toxicity, as a critical determinant for informing choice of treatment, particularly in situations where the MIC may not be achieved.

## Materials and methods

### Chemicals, reagents and antibodies

Ciprofloxacin, moxifloxacin, tobramycin and gentamycin were purchased from Merck. The α-FLAG antibody was purchased from Merck and α-GAPDH from Abcam. The pcrV antibody Mab 166 was purchased from Creative Biolabs (New York, USA). The pOPINF *E. coli* expression vector was purchased from Addgene. The pUCPT20 encoding GFP for constitutive gene expression in *P. aeruginosa* was also purchased from Addgene. ExoU with a C-terminal 6xHistidine tag was also cloned into pUCPT20 in place of the GFP gene and transformed into PA103 ΔUT where indicated. LIVE/DEAD assay reagents were purchased from Invitrogen. LDH assay reagents were purchased from Thermofisher. Arachidonoyl thio-PC was purchased from Cayman Chemical (Michigan, USA). PIP_2_ was purchased from Avanti Polar Lipids (Alabama, USA). Bovine ubiquitin and 5,5’-dithiobis- (2-nitrobenzoic acid) (DNTB) were purchased from Sigma-Aldrich. ExoU inhibitor Compound B was purchased from Chem Bridge (California, USA).

### Bacterial strains used in this study

The clinical isolate strain of *P. aeruginosa*, PA103, and a PA103 ExoU and ExoT knock out mutant (PA103 ΔUT) [28] were a gift from Professor Dara Frank (Medical College of Wisconsin). PA76026 is a clinically genotyped and phenotyped ExoS expressing strain that was obtained from the University of Liverpool which houses isolates of the Microbiology Ophthalmic group. The pUCPT20 encoding either GFP or ExoU with a C-terminal 6xHistidine tag were transformed into PA103 and PA76026 by electroporation with 300 μg/ml carbenicillin employed as the selection marker. Recombinant ExoU with an N-terminal 6xHistidine tag was expressed in C43 *E. coli*, and purified from *E. coli* as previously described [29].

### Detection of *P. aeruginosa* colony forming units

Cultures of *P. aeruginosa* in LB broth, with and without indicated antimicrobials, were centrifuged, resuspended in 1 mL of PBS, serially diluted and then incubated on agar plates overnight at 37 °C prior to counting of colony forming units (CFUs). For detection of extracellular bacteria after 24 hours HCE-T cell infection, cell culture medium was removed, centrifuged at 5000 x *g* and bacteria resuspended and serially diluted in PBS followed by plating onto LB agar. After overnight incubation CFU/mL were quantified. For detection of intracellular bacteria, 200 μg/mL of gentamycin was added to the culture medium for 2 hours to eliminate extracellular bacteria. Gentamycin was removed and the HCE-T cells were washed with PBS. HCE-T cells were collected by scrapping in buffer containing 20 mM Tris (pH 7.4), 100 mM NaCl and 0.05% (v/v) Triton. Suspensions were briefly sonicated with two pulses of 3 seconds at an amplitude of 5, serially diluted in PBS, spread on to agar plates, incubated overnight at 37 ºC and finally colonies were counted.

### Western blotting

Bacteria were isolated by centrifugation at 5000 x *g* for 5 minutes. After resuspension in lysis buffer (50 mM Tris-HCl (pH 7.4), 1% (v/v) NP-40, 0.1% (v/v) SDS, 100 mM NaCl, 1 mM DTT, 10% glycerol and cOmplete protease inhibitor cocktail (Roche)), bacteria were briefly sonicated on ice and then centrifuged at 16,000 x *g* prior to protein quantification using the Bradford assay. Samples were boiled for 5 minutes in sample buffer (50 mM Tris-HCl (pH 6.8), 1% (v/v) SDS, 10% (v/v) glycerol, 0.01% (w/v) bromophenol blue, and 10 mM DTT). Subsequently, 80 μg of total protein for each sample was resolved by SDS-PAGE prior to transfer to nitrocellulose membranes (Bio-Rad). Membranes were blocked in Tris-buffered saline + 0.1% (v/v) Tween 20 (TBS-T) in 5% (w/v) milk (pH 7.4) followed by incubation with indicated primary antibodies overnight. Proteins were detected using appropriate secondary HRP-conjugated antibodies and enhanced chemiluminesence reagent (Bio-Rad). Band intensities were quantified using ImageJ software.

### qRT PCR

Bacteria were sub-cultured at an OD_600_ of 0.1and then grown in a shaker incubator at 37ºC for 16 hours in the presence of indicated antimicrobial agent. Cells were collected by centrifugation and lysed in RLT buffer (Qiagen) according to the manufacture’s instructions. The mRNA was extracted using an RNA extraction kit (Qiagen). Complete cDNA was generated from total RNA using GoScript Reverse Transcription system (Promega), using 1 μg RNA per reaction and 0.5 μg of Random primer. qPCR was performed in triplicate using the Comparative Ct (ΔΔCt) method on an Applied Biosystems (AB) StepOnePlus machine, a Power SYBR Green PCR Master Mix (Thermo Scientific) and the following primer pairs. Expression levels were normalised to AmpC mRNA.

ExoU: left 5′-AGAACGGAGTCACCGAGCTA and right 5′-CGAGCAGCGAAATAAGATCC.

ExoS: left 5′-ATGTCAGCGGGATATCGAAC and right 5′-CCTCAGGCGTACATCCTGTT.

pcrV: left 5′-TGATCCAGTCGCAGATCAAC and right ATCCTTGATCGACAGCTTGC.

ExsA: left 5′-TTGAGTGAAGTCGAGCGTTG and right 5′-TCCATGAATAGCTGCAGACG.

AmpC: left 5′-ACCCATCGCGGTTACTACAA and right 5′-GTGGAACCGGTCTTGTTCAG.

### In vitro PLA2 assay

Recombinant (from *E. coli*) and endogenous (secreted from PA103) ExoU sn-2 directed phospholipase activity was detected using an adapted Caymen chemical cPLA2 assay kit in a 96-well plate format, as previously described [29]. Assay conditions contained 1mM ATPC, 1 μM PIP2, 25 μM mono ubiquitin, 2% DMSO (v/v) and 1.25 mM 5,5-dithio-bis-(2-nitrobenzoic acid) (DTNB) in a final volume of 50 μL. For detection of recombinant ExoU phospholipase activity, 100 nM of ExoU was added to initiate substrate hydrolysis. For detection of endogenous ExoU secreted from PA103, 25 μL of culture medium was used. The absorbance at 405 nm (A405) was measured and background subtracted (substrate and DTNB alone) at 2-minute increments over 3 hours (for recombinant ExoU) and 24 hours (for endogenous ExoU). For recombinant ExoU, substrate hydrolysis was calculated using the equation A405/10.00 × 0.05 ml/number of nanomoles of ExoU for the 96-well plate format and A405/10.00 × 0.01 ml/number of nanomoles of ExoU. Endogenous ExoU activity was calculated as a percentage of DMSO controls with no antimicrobial treatment.

### HCE-T scratch and infection assay

HCE-T cells were analysed using a scratch and infection assay as previously described [29]. Briefly, HCE-T cells were cultured to fully confluent monolayers in 24-well plates. Two parallel scratches were applied across the diameter of the wells with a pipette tip. PA103 and PA76026 were added at a multiplicity of infection (MOI) of 2.5 with the indicated antimicrobial or DMSO (0.01% v/v) controls.

### Fluorescence microscopy

Scratched and infected HCE-T cells with or without antimicrobials were incubated at 37 °C in 5% CO_2_ for 24 hours before analysis by florescent microscopy, employing Live/Dead staining (Invitrogen), to differentiate and visualise viable and dead/dying cells. Culture medium was removed from the infected HCE-T cells and washed with 1 ml of PBS three times. Fresh medium containing 5 μM of both Calcein (Ex/Em 494/517 nm) and Ethidium homodimer-1(Ex/Em 528/617 nm). Fluorescence microscopy was also used to detect GFP labelled internalised PA76026. HCE-T cells were infected with PA76206 transformed with pUCPT20 GFP at a MOI of 2.5, in the presence of indicated antimicrobial for 24 hours. Cells were washed with PBS and then visualised (Ex/Em 395/509). Images of the scratched HCE-T cells were obtained on either an Apotome Zeiss Axio Observer or a Nikon Eclipse TiE.

### LDH assays

As an indicator of cell lysis, lactate dehydrogenase (LDH) release was measured using the Pierce LDH Cytotoxicity Assay Kit (Thermo Scientific) according to the manufacturer’s instructions. Culture medium (50 μL) of HCE-T cells was assayed 24 hours after initial infection with the indicated strains of *P. aeruginosa* in the presence of indicated antimicrobial (0.1% DMSO v/v). The results were reported as percent LDH release normalised to a positive control (according to manufactures instructions), which gave the maximum amount of observable cell lysis in an appropriate detectable range of absorbance.

## Results

### Analysis of fluoroquinolones and aminoglycosides on *P. aeruginosa* growth

Our aim was to analyse the effects of antimicrobials on T3SS virulence factor expression below their respective minimal inhibitory concentrations (MICs). For this purpose, we first established the MIC_50_ (concentration required for 50% bacterial growth inhibition) for two fluoroquinolones (moxifloxacin and ciprofloxacin) and two aminoglycosides (tobramycin and gentamycin) on the growth of the *P. aeruginosa* strains PA103 and PA76026. PA103 expresses ExoU whereas PA76026 expresses ExoS. The amount of growth after 16 h in the presence of the antimicrobials was determined, and this revealed that both *P. aeruginosa* strains were more susceptible to inhibition by the fluoroquinolones moxifloxacin and ciprofloxacin, than the aminoglycosides tobramycin or gentamycin (Figure 1A). We established that the MIC_50_ for ciprofloxacin, moxifloxacin, tobramycin and gentamycin were 0.5, 2, 6, and 8 μM for PA103 and 1, 2.5, 6 and 8 μM for PA76026.

**Figure 1:**
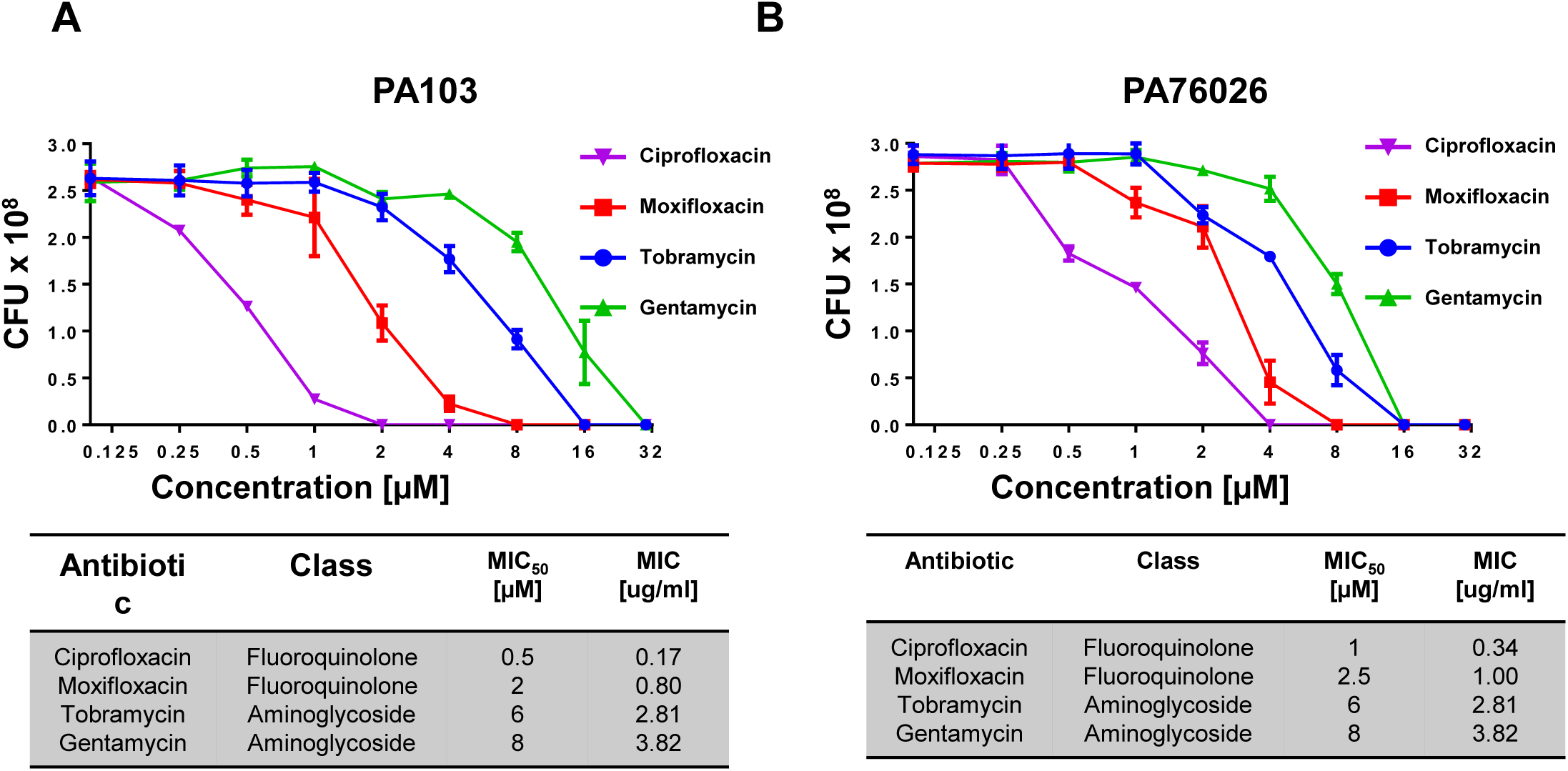
Antibiotic minimal inhibitory concentrations for ExoU expressing PA103 and ExoS expressing PA76026 strains of *P. aeruginosa*. The antibiotic at 50% minimal inhibitory concentration (MIC_50_) for ExoU expressing PA103 (A) and ExoS expressing PA76026 (B) strains of *P. aeruginosa* were determined by measuring absorbance reading at OD_600_ nm to assess bacterial growth, after cultures were incubated with varying concentrations of specified antibiotic in 96-well plates. The antibiotic type and their minimal inhibitors concentrations for growth of PA103 and PA76026 are displayed.

### Effects of antimicrobials on PcrV expression in *P. aeruginosa*

We next used western blotting to detect changes in expression of the essential T3SS needle tip component, PcrV [9], in PA103 and PA76026 after 16 hours incubation with antimicrobials at their respective MIC_50_ (Figure 2A). Although less potent than ciprofloxacin and moxifloxacin, the aminoglycoside tobramycin, at 6 μM, caused a sharp reduction in PcrV produced by both PA103 (74.0% reduction) and PA76026 (50.5% reduction) (p = 0.001 μM and 0.003 μM). Gentamycin, also an aminoglycoside, did not detectably alter PcrV expression for either cell line. The fluoroquinolone moxifloxacin, however, caused a statistically significant increase in the relative abundance of PcrV in PA103 (81.8% increase, p = 0.004), whereas ciprofloxacin, caused a similar increase in PcrV expression in PA76026 (57.0%, p = 0.003) (Figure 2A).

**Figure 2:**
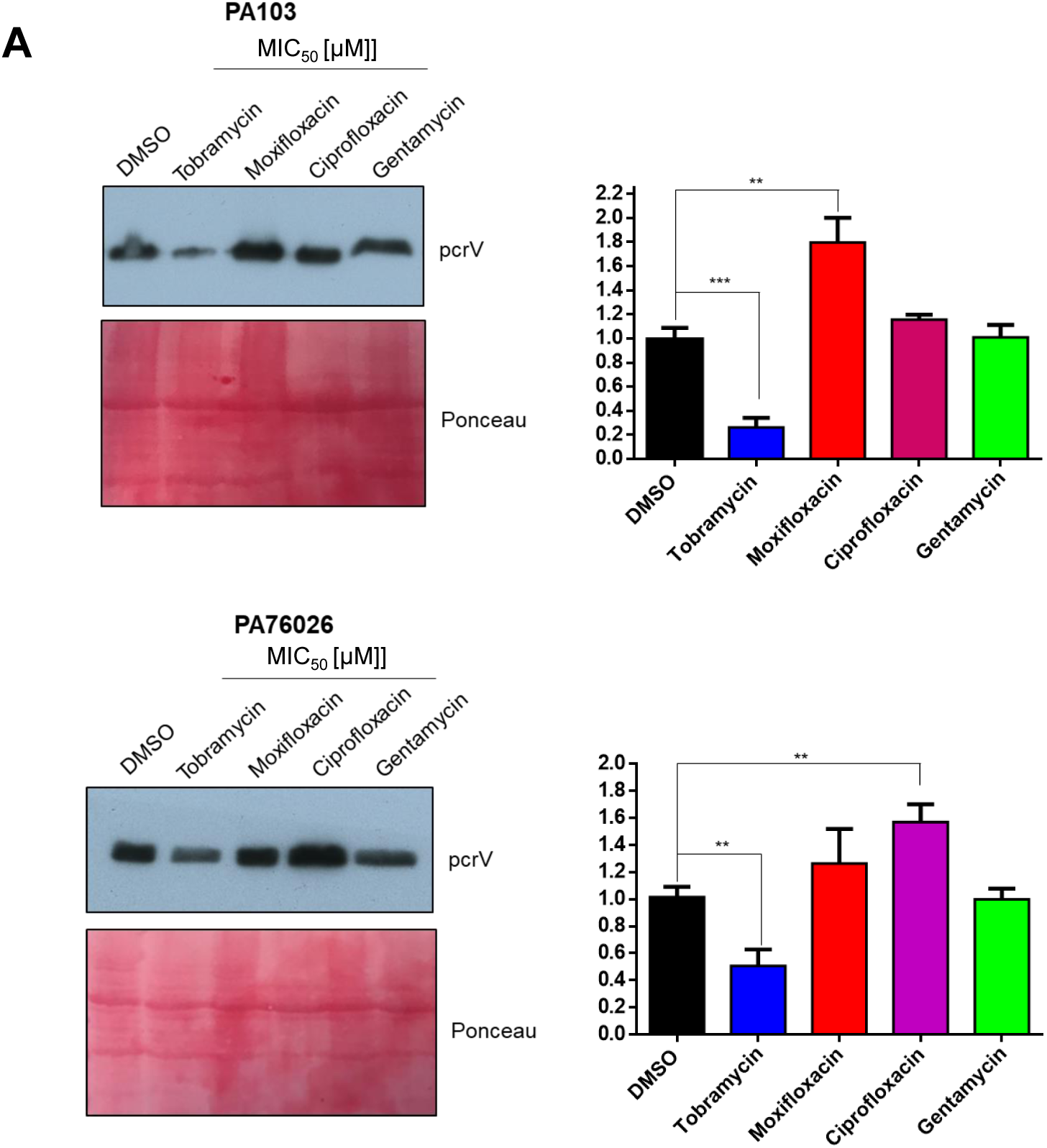
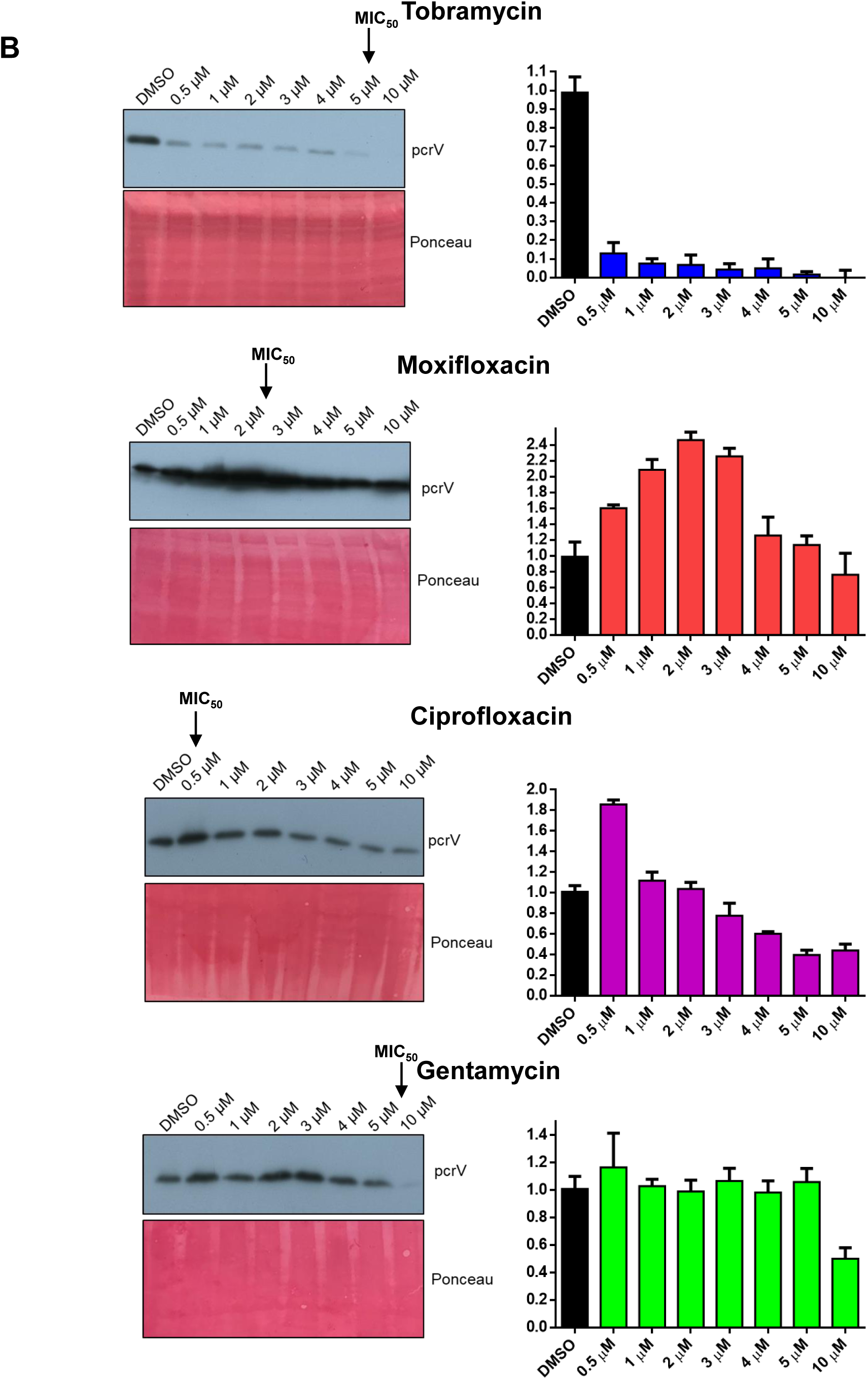
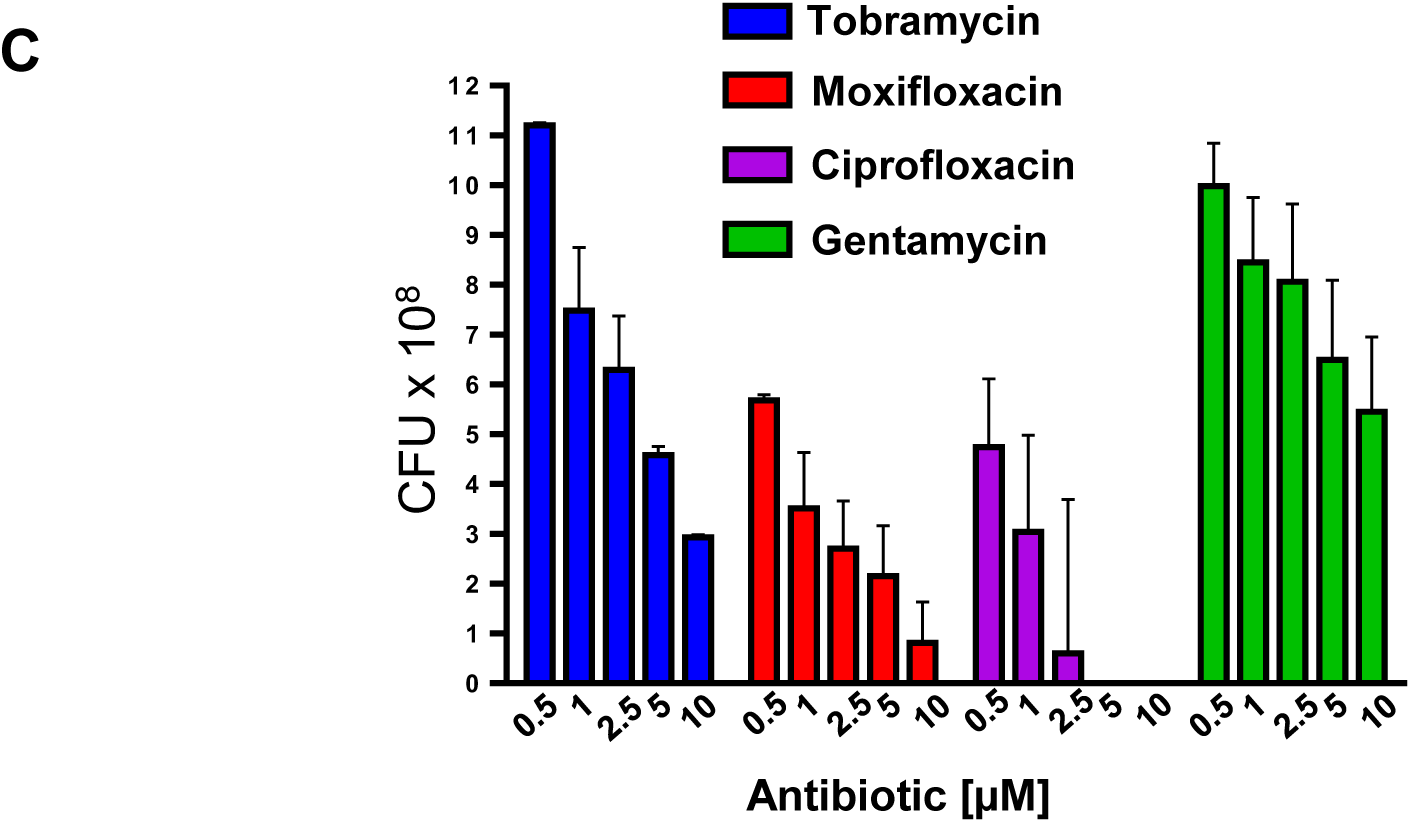
Tobramycin is less potent then moxifloxacin and ciprofloxacin but reduces PcrV expression in *P. aeruginosa*. (A) Expression of PcrV in PA103 and PA76026 after 16 hours incubation with indicated antibiotic at the MIC_50_, determined by western blotting. Relative band intensities were calculated using imageJ software from 3 independent experiments. T-tests were used to determine statistically significant difference in relative PcrV expression levels. (B) The effect of antibiotic, at increasing concentrations on PcrV expression in PA103 was determined by western blotting. Relative band intensities were determined from 3 independent experiments. (C) The number of viable PA103 CFUs were detected 16 hours after incubation with varying concentrations of indicated antibiotic.

To evaluate changes in PcrV expression after antimicrobial exposure in more detail, PA103 was exposed to antimicrobials at varying concentrations prior to western blotting (Figure 2B) and colony forming unit (CFU) analysis (Figure 2C). Even at 0.5 μM (4.2% of the MIC), tobramycin caused a stark reduction in detectable PcrV (Figure 2B). In contrast, PcrV increased at moxifloxacin concentrations between 0.5 and 3 μM. Only at concentrations above the MIC of moxifloxacin did the relative abundance of PcrV return to basal levels. Ciprofloxacin, which is a more potent antimicrobial than moxifloxacin (MIC_50_ of 0.5 μM), caused an increase in PcrV expression at 0.5 μM, but induced a concomitant reduction in PcrV at concentrations >4 μM. The aminoglycoside gentamycin only induced loss of PcrV expression at concentrations above 10 μM (Figure 2B). Analysis of viable PA103 CFUs following antimicrobial treatment (Figure 2C) largely reflected our previous observations for cell growth (Figure 1A), with the two fluoroquinolone compounds proving the most toxic and the two aminoglycosides only modestly inhibiting cell viability at the highest tested concentrations. The high CFU count for tobramycin treated PA103 is therefore in stark contrast to induced loss of PcrV protein (Figure 2B).

### Analysis of T3SS-related gene transcription in *P. aeruginosa* in response to antimicrobial exposure

To further investigate how antimicrobials impact expression of the *P. aeruginosa* T3SS complex and associated cytotoxins at the transcriptional level, we used qRT-PCR (Real-Time Quantitative Reverse Transcription PCR) to detect changes in mRNA levels for *exoU* (for PA103), *exoS* (for PA76026), *pcrV* and the key T3SS activating transcription factor, *exsA*. EGTA, which has previously been shown to increase *exsA* transcription in *P. aeruginosa*, was used as a positive control to induce T3SS expression [13, 30]. After incubation with 2 mM EGTA for 16 h, there was predictably an increase in *exoU* (PA103), *exoS* (PA76026), PcrV and ExsA mRNA in the *P. aeruginosa* clinical isolates (Figure 3A and B). With 1 μM of tobramycin, we observed a 0.6 fold decrease in *exoU* mRNA in PA103, relative to DMSO treated controls (Figure 3A, left), whilst mRNA levels of *pcrV* and *exsA* were unaffected. With 1 μM of moxifloxacin, *exoU* and *pcrV* mRNA levels did not significantly change, although, we did observe a 1.4-fold increase in *exsA* mRNA (Figure 3A, left). In the presence of 5 μM of tobramycin (83.3% of the MIC_50_), ExoU expression decreased further (0.3 fold), but *pcrV* and *exsA* mRNA was consistent with DMSO treatment (Figure 3A, right). Increasing the concentration of moxifloxacin to 2 μM (0.5 μM below the 2.5 μM MIC_50_) resulted in pronounced increases in *pcrV* (2.7 fold) and *exsA* (1.8 fold) mRNA (compared to DMSO), with exoU remaining relatively unchanged (Figure 3A, right). The observed increase in *pcrV* and *exsA* expression at the transcriptional level is consistent with the changes we previously observed at the protein level in moxifloxacin treated PA103 cells (Figure 2).

**Figure 3:**
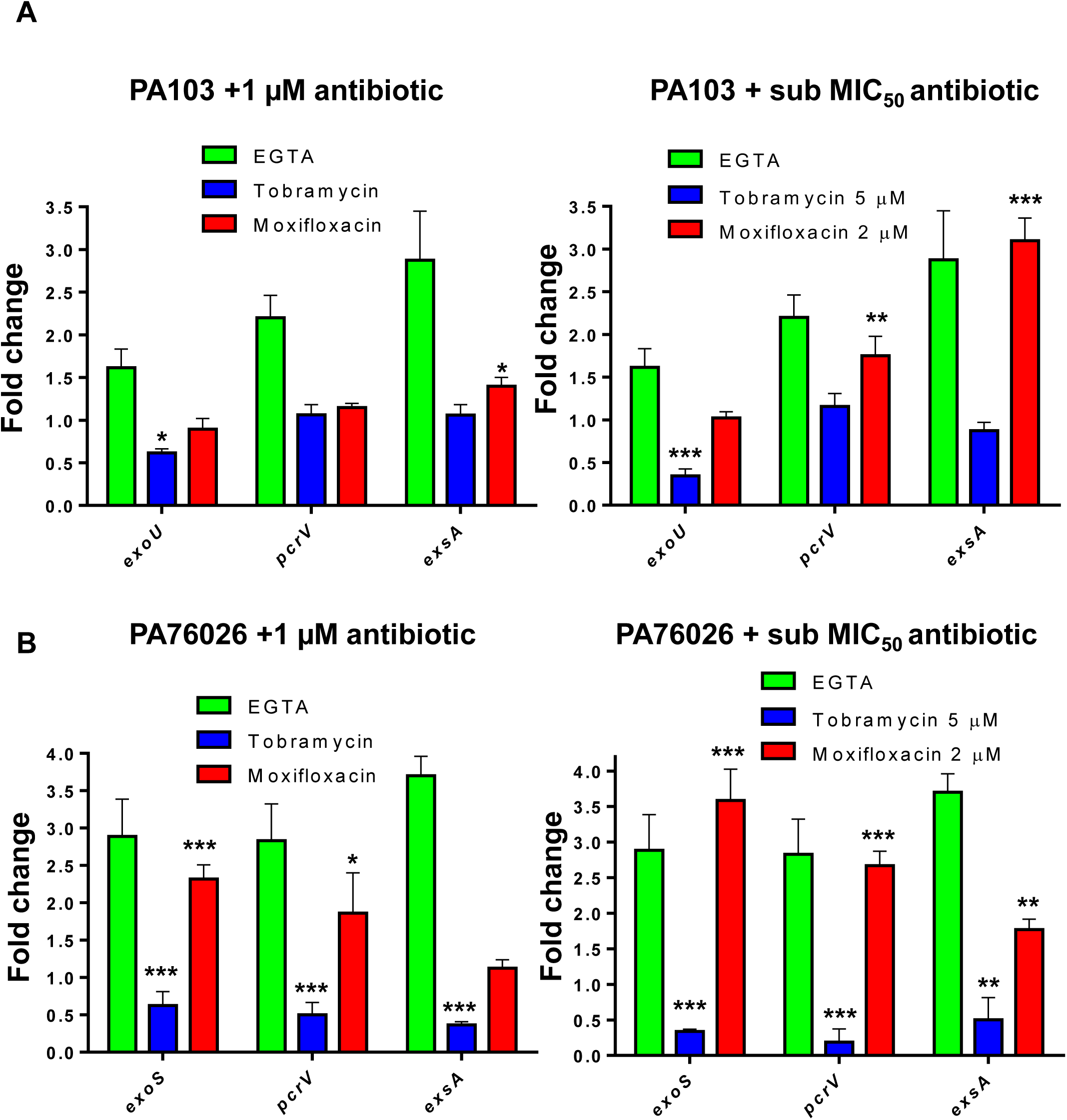
Impact of tobramycin and moxifloxacin on T3SS gene expression in PA103 and PA76426. PA103 (A) and PA76026 (B) were incubated for 16 hours in the presence of either 1 μM (left) or the antibiotic MIC_50_ (right) prior to RT-qPCR analysis to detect relative mRNA levels of T3SS associated genes. Incubation with 2 mM of EGTA served as the positive control for T3SS induction. Values are means ± SD from 3 independent experiments; *p < 0.05; **p < 0.01; ***p < 0.001.

For PA76026, we observed downregulation of all three T3SS-associated proteins in the presence of 1 μM tobramycin, (Figure 3B). *exoS* mRNA transcription decreased by 0.6 fold, *pcrV* by 0.5 fold and *exsA* by 0.4 fold in PA76026 treated (Figure 3B, left). Conversely, in the presence of 1 μM moxifloxacin we observed upregulation of *exoS* and *pcrV* (2.3% and 1.9 fold, respectively) whilst *exsA* mRNA levels were unaffected (Figure 3B, left). Similar to 1 μM tobramycin exposure, we detected decreases in mRNA of *exoS* (0.3 fold), *pcrV* (0.2 fold) and *exsA* (0.5 fold) when the concentration was increased to 5 μM (Figure 3B, right). With 2 μM moxifloxacin, we also observed a sharp increase in mRNA for *exoS* (3.6 fold), *pcrV* (2.7 fold) and even *exsA* (1.8 fold).

### Effect of antibiotics on ExoU secretion by PA103

Since tobramycin reduced T3SS expression, as judged by a diminished PcrV protein signal (Figure 2), and inhibited ExoU expression at the transcriptional level, we next probed for accompanying modulation in ExoU secretion by PA103 cells. To this end, PA103 was incubated with either tobramycin, moxifloxacin, ciprofloxacin and gentamycin (at the respective MIC_50_) for 16 hours after which point, the culture medium (containing secreted ExoU) was analysed using a phospholipase assay (Figure 4A). In the presence of tobramycin, but not moxifloxacin, ciprofloxacin or gentamycin, there was a decrease in ExoU phospholipase activity detected in the culture medium. Importantly, none of these antimicrobials inhibited the enzymatic activity of recombinant His-tagged ExoU expressed in and purified from *E. coli* (supplementary figure 1), indicating that tobramycin prevented ExoU production and/or secretion rather than having a direct inhibitory effect on ExoU’s activity. To eliminate the possibility that the observed differences in PLA_2_ activity was a consequence of antibiotic toxicity as opposed to a specific reduction in ExoU production and/or secretion, we also verified that CFU counts for all treated conditions were comparable (Figure 4B).

**Figure 4:**
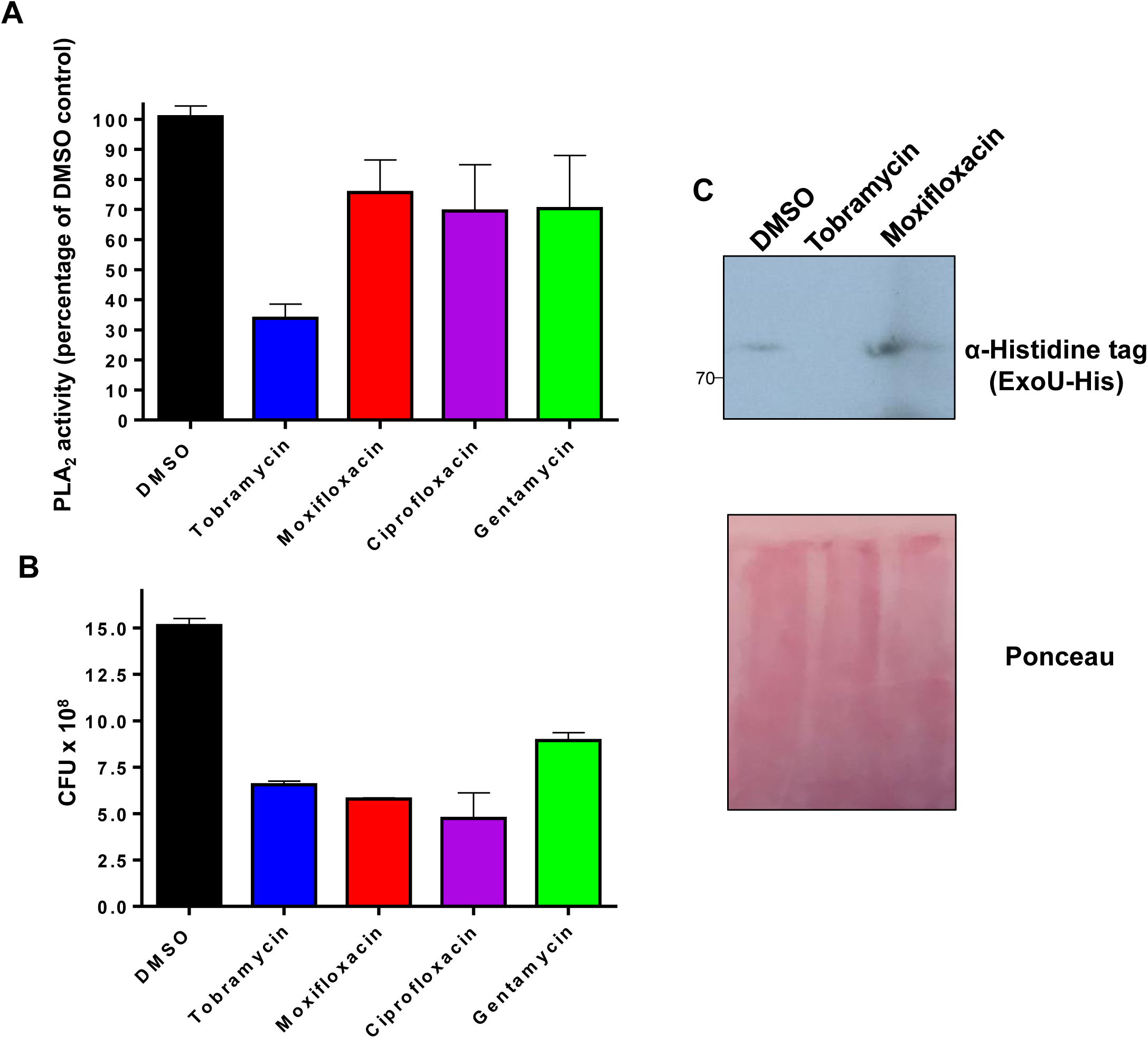
Tobramycin prevents secretion of ExoU. (A) PA103 was incubated for 16 hours in the presence of indicated antibiotic at their respective MIC_50_.The cleared bacterial culture medium was then analysed employing a PLA_2_ phospholipase assay in order to detect ExoU activity, with ubiquitin (20 μM) and PIP_2_ (1 μM) present as ExoU activating co-factors. Results are shown as percentage ExoU activity after 24 hours relative to ExoU secreted from DMSO control treated PA103. (B) The number of PA103 CFUs were detected after 16 hour MIC_50_ antibiotic exsposure. (C) PA103 ΔUT: pUCP ExoU-His was incubated with DMSO, tobramycin and moxifloxacin for 16 hours. The cleared culture medium was then analysed by western blotting, employing an anti-6xhistidine primary antibody, in order to detect secreted His-tagged ExoU.

As (to our knowledge) there are currently no commercially available ExoU antibodies, we transformed an ExoU and ExoT knock out strain of PA103 (PA103 ΔUT) with a pUCPT20 plasmid encoding ExoU modified with a C-terminal 6x histidine tag to serve as an artificial antigen for immunogenic detection. We next quantified secreted ExoU in the culture medium by western blotting (Figure 4C). When PA103 ΔUT pUCPT20-ExoU-His was incubated overnight with 0.01% (v/v) DMSO, ExoU was readily detected in the medium (Figure 4C). In the presence of 6 μM of tobramycin however, secreted ExoU was essentially eliminated. Interestingly, 2.5 μM of moxifloxacin, resulted in an increase in ExoU-in the culture medium (Figure 4C).

### Effects of antimicrobials on PA103 cytotoxicity in a wound healing infection model

In a previous study, we developed a corneal epithelial HCE-T cell scratch and infection assay to evaluate inhibitors of ExoU as an *in vitro* model of disease [29]. Infection and ExoU cytotoxicity is established along the border of the scratch, preventing healing and leading to a widening of the wound, which can be observed by fluorescence microscopy, while ExoU-specific cytotoxicity can simultaneously be indirectly estimated by LDH assays. We set out to determine how antimicrobials (at varying concentrations) might influence acute ExoU-driven cytotoxicity after infection of HCE-T cells with PA103, using LIVE/DEAD fluorescence microscopy analysis to observe HCE-T cell viability and wound healing (Figure 5A) in addition to quantifying PA103 CFUs in the culture medium following treatment (Figure 5B). Finally, we used an LDH assay to determine the degree of cell lysis, the data for each condition presented as a proportion of the antimicrobials MIC_50_ (Figure 5C).

**Figure 5:**
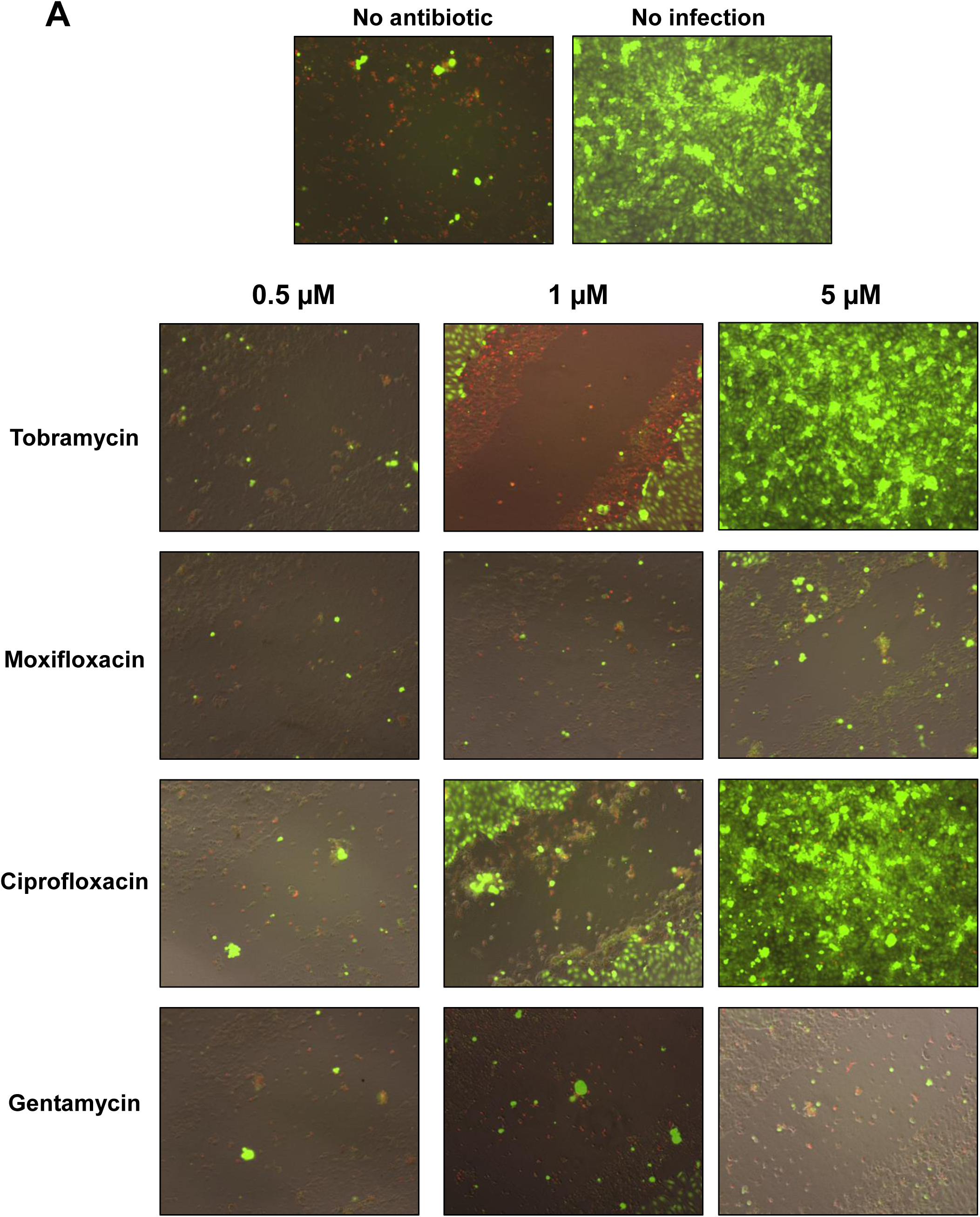
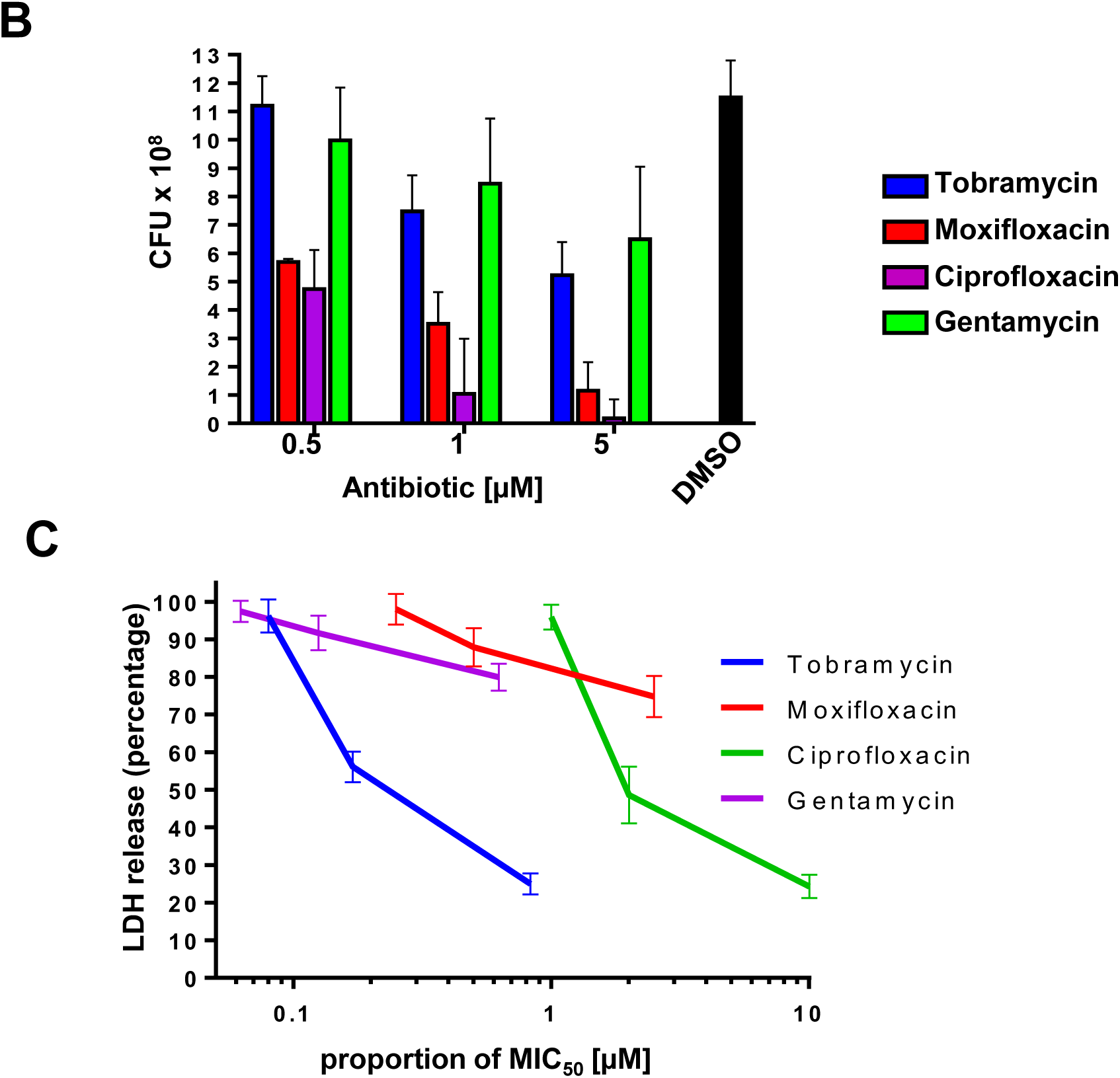
Tobramycin prevents ExoU mediated cytotoxicity in a HCE-T scratch and infection assay. (A) Live/Dead fluorescence microscopy analysis of scratched HCE-T cells 24 h post infection with ExoU expressing PA103 in the presence of varying concentrations of indicated antibiotic. (B) Number of PA103 CFUs in the cell culture medium 24 h after infection. (C) Percentage LDH release from infected HCE-T cells as a marker of ExoU induced cell lysis in the presence of varying doses of antibiotic, expressed as a proportion of the antibiotics respective MIC_50_.

After scratched HCE-T cells were incubated for 24 hours with PA103 without antimicrobials (DMSO 0.01% v/v), almost all of the cells had succumb to infection, (figure 5A, top left). Reciprocally, when no PA103 was present we observed complete wound closure (Figure 5A, top right). Tobramycin was able to partially mitigate cytotoxicity at 1 μM (6 times lower than the MIC_50_), despite extensive bacterial expansion (Figure 5B). This was also reflected in a reduction of LDH release (49.3%) compared to DMSO (Figure 5C). Exposure to 5 μM tobramycin resulted in complete wound closure (Figure 5A) and a further reduction in LDH release (Figure 5C), signifying an abrogation of PA103 cytotoxicity. Interestingly, despite inducing a significant reduction in CFU compared to DMSO treated control samples, treatment with 5 μM tobramycin was consistently less bactericidal than similar concentrations of moxifloxacin and ciprofloxacin (Figure 5B). Moxifloxacin at all concentrations tested was ineffective at reducing cytotoxicity (Figure 5A and 5C), seemingly in contradiction to its higher bactericidal potential in comparison to tobramycin (Figure 5B). Ciprofloxacin was not effective in preventing cytotoxicity at its MIC_50_ (0.5 μM). Above the MIC_50_ (1 μM), ciprofloxacin was able to partially rescue scratched HCE-T cells from PA103 mediated lysis (Figure 5A and 5C), which coincided with extensive bacterial death (Figure 5B). At 5 μM of ciprofloxacin we observed almost total bacterial clearance and wound closure. Gentamycin was not effective in preventing cytotoxicity, although, all concentrations tested were below gentamycin’s MIC_50_ of 8 μM.

### Effect of antimicrobials on ExoS cytotoxicity and intracellular survival of PA76026

We next sought to determine whether tobramycin, moxifloxacin, ciprofloxacin or gentamycin could prevent T3SS mediated cytotoxicity from the ExoS expressing strain of *P. aeruginosa*, PA76026. In our scratch and infection assay, ExoS activity caused cell rounding along the site of the initial scratch after 6 hours (supplementary figure 2). Cell rounding in this manner due to ExoS activity has been reported previously (Frank 2018). In the absence of an antimicrobial, cell death occurred after 24 hours due to bacterial expansion, which overwhelmed the culture medium and therefore may not be attributed to ExoS action alone (supplementary figure 2). With antimicrobial at their respective MIC_50_, the bacterial load was controlled and we could observe the effects of ExoS toxicity in scratched HCE-T cells over 24 hours (Figure 6). We used LIVE/DEAD fluorescence microscopy to observe cell viability and morphological changes. We also determined both extra- and intracellular bacterial CFUs following 24 hours of infection, as ExoS activity has previously been reported to promote intracellular survival of *P. aeruginosa* [13, 31, 32].

**Figure 6:**
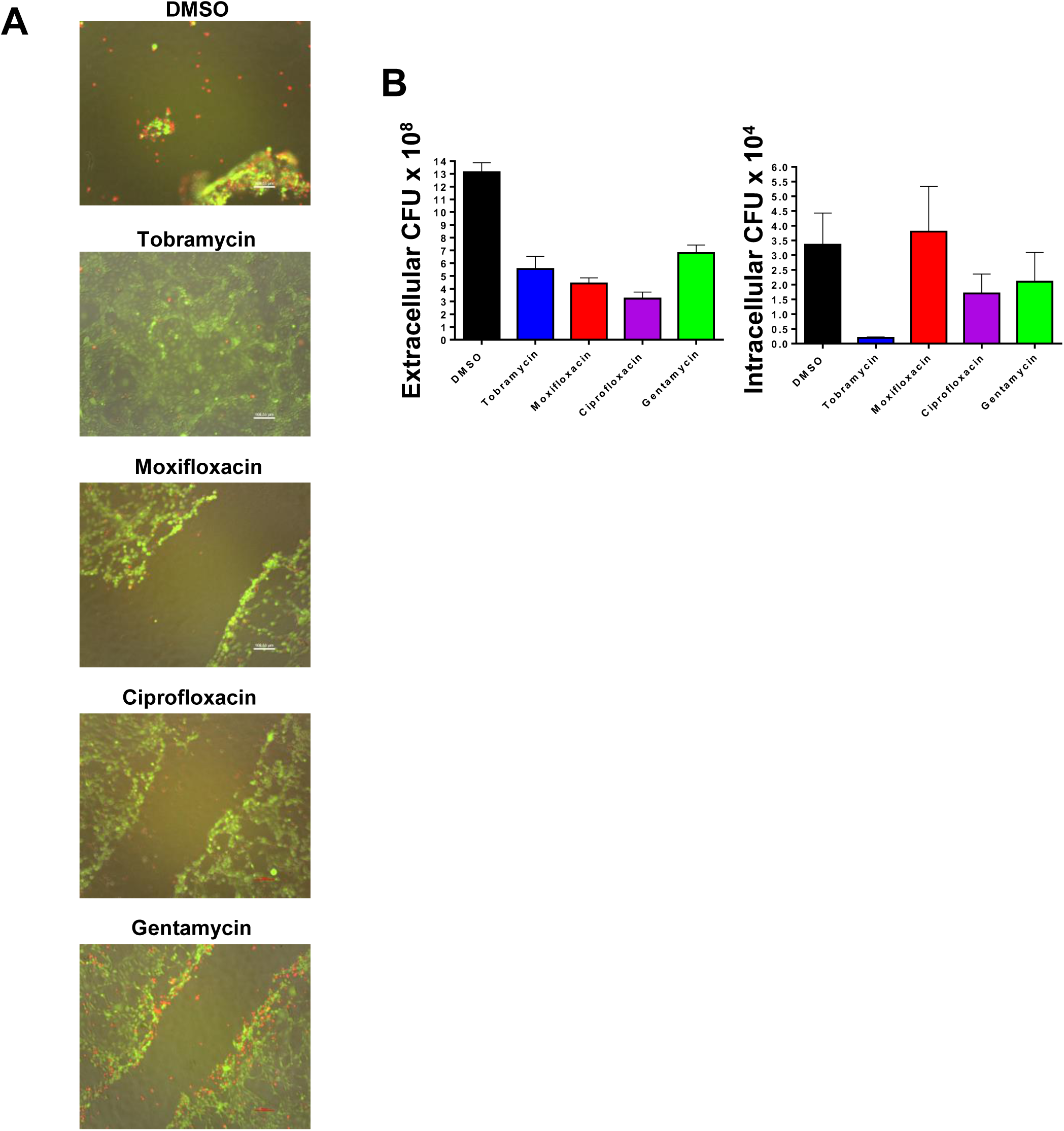
Tobramycin reduces intracellular survival of ExoS expressing PA76026 at MIC_50_ during HCE-T cell infection. (A) Live/Dead fluorescence microscopy analysis of scratched HCE-T cells 24 h post infection with ExoS expressing PA76026 in the presence indicated antibiotic at the MIC_50_. (B) Number of extracellular (left) and intracellular (right) CFUs of PA76026,with different antibiotic treatments 24 hour post infection, were detected.24 h after infection.

Employing Live/Dead fluorescence microscopy we observed that tobramycin, at the MIC_50_, abolished PA76026 mediated cytotoxicity, resulting in complete wound closure (Figure 6A). Despite similar extracellular quantities of bacteria in all treatment conditions, tobramycin induced a massive reduction in intracellular CFUs (Figure 6B). Conversely, moxifloxacin, ciprofloxacin and gentamycin, at their respective MIC_50_, did not facilitate wound healing and we observed numerous rounded cells along the border of the scratch (Figure 6A). This correlated with an increase in intracellular CFUs relative to tobramycin treatment (Figure 6B). In addition, we transformed our PA76026 strain with a GFP encoding pUCPT20 plasmid so that intracellular bacteria could be observed by fluorescence microscopy (supplementary figure 3). Interestingly, we could detect more intracellular PA76026 in moxifloxacin treated HCE-T cells than any other condition.

## Discussion

ExoU and ExoS expressing strains of *P. aeruginosa* are related to poorest prognosis in pneumonia and contact lens associated keratitis [2, 10, 33]. The current treatment for *P. aeruginosa* keratitis is the prescription of multiple antibiotics, which must be introduced rapidly following the onset of symptoms to minimise corneal damage [34]. This approach often results in in corneal toxicity and selection for antibiotic-resistance [35], leading to failure of treatment. Therefore, a better understanding of the effects of antimicrobials on *P. aeruginosa* virulence will be critical for developing improved therapeutic strategies.

The aminoglycosides, tobramycin, at concentrations well below the calculated MIC_50_ caused a sharp reduction in the T3SS secretion apparatus protein, PcrV (Figure 2A and B) in ExoU expressing PA103 cells, which was accompanied by reduced ExoU secretion and phospholipase activity (Figure 4). We also detected a tobramycin-dependent loss of *exoU* mRNA signal, which could partially explain this latter observation. The other aminoglycosides investigated in this study, gentamycin also reduced PcrV expression, but only at concentrations above the MIC_50_ (Figure 2B). Interestingly, PA103 expression of *pcrV* and the T3SS activating transcription factor, *exsA*, were unaffected by tobramycin at the transcriptional level at the concentrations used in this study. In contrast, a detectable loss in mRNA signal for *exoS, pcrV* and *exsA* was observed in PA76026 cells exposed to tobramycin. Aminoglycosides inhibit bacterial protein synthesis [25], and we hypothesised that is the dominant mode of action to explain the reduced expression of PcrV, ExoS and ExoU that we observed for the two clinical strains of *P. aeruginosa* used in this study.

Fluoroquinolones, such as moxifloxacin, function by inhibiting bacterial DNA replication by targeting DNA topoisomerase and DNA-gyrase [23]. Here, we observed that moxifloxacin (at sublethal concentrations) increased total *exsA* and *pcrV* mRNA in both isolates, and *exoS* in PA76026 cells (but not *exoU* in PA103), which collectively suggested a general upregulation of T3SS expression that is comparable to the established T3SS inducing agent, EGTA. This also correlated with a concentration dependent increase in PcrV protein for PA103 challenged with moxifloxacin below the MIC_50_. This highlights the concerning possibility that targeting *P. aeruginosa* with fluoroquinolones, particularly at sub-lethal concentrations, might provoke T3SS expression and enhance acute pathogenicity.

### Prevention of *P. aeruginosa* toxicity by tobramycin in a wound healing model

Neither moxifloxacin or gentamycin were effective at preventing *P. aeruginosa* mediated cytotoxicity at or below 5 μM in a corneal HCE-T scratch and infection model (Figure 5). PA103 infected corneal cells were viable with ciprofloxacin application at five times the calculated MIC, but we believe this is primarily due to extensive bacterial clearance by the highly bactericidal compound (Figure 5A and B). Interestingly, tobramycin was relatively ineffective at reducing bacterial loads, but afforded potent protection of HCE-T towards infection and cytotoxicity, which we partially attribute to a depletion in TS33 mediated toxicity and ExoU secretion. A previous study revealed that tobramycin was effective at reducing acute cytotoxic damage and could decrease neutrophil extracellular trap (NET) formation in a mouse keratitis model of *P. aeruginosa* infection [27]. Although the authors could not account as to how tobramycin mitigated NET formation, our results might offer insight. Proinflammatory signalling in the host, induced by T3SS effectors, has been shown to potentiate deleterious effects of neutrophil infiltration leading to tissue damage [36, 37]. Antimicrobials such as amoxicillin have been shown to increase NET formation [38], leading to exacerbated tissue damage, whereas gentamycin was shown to reduce NET formation [39]. This suggests that particular antimicrobials may fail in certain therapeutic circumstances, whereas other antimicrobial classes could be of benefit.

### Prevention of ExoS toxicity and intracellular survival of *P. aeruginosa* by tobramycin

Retention of an intracellular phenotype, indicative of ExoS activity, was observed for PA76026 in the presence of moxifloxacin, ciprofloxacin and gentamycin, as evidenced by HCE-T cell rounding along the border of the scratch, as well as the detection of viable intracellular CFU (Figure 6A and B). This was particularly apparent in moxifloxacin containing assays, as intracellular bacterial CFU counts were comparable to control experiments. Furthermore, we detected enhanced GFP-expressing intracellular PA76026 (supplementary Figure 3). Our initial findings indicating a moxifloxacin dependent upregulate the T3SS system, including the pro-intracellular survival virulence factor, ExoS, may partially explain these latter observations. In contrast, tobramycin significantly ablated intracellular populations of PA76026, which also manifest as complete wound healing in our scratch assay. The apparent discrepancy in the actions of both compounds when applied at sub-lethal concentrations (for *P. aeruginosa*), in regards to antimicrobial potential, is likely partially a consequence of tobramycin impeding T3SS mediated cytotoxicity. However, given that the mode of action of tobramycin and aminoglycosides is to block bacterial protein synthesis by binding directly to A-site on the 16S ribosomal RNA of the 30S ribosome, it remains to be seen how specific this inhibition of T3SS secretory apparatus is. Undoubtedly however, interference of T3SS, and thus secretion of ‘pro-intracellular survival’ virulence factors such as ExoS, is likely a major contributary factor in the loss of viable invasive PA76026 cells, at concentrations of tobramycin determined to be only minimally bactericidal in isolation. Although gentamycin is also an aminoglycoside, it was only able to reduce PcrV expression at concentrations exceeding the MIC_50_ (Figure 2B), which we believe is why gentamycin offered limit protection in wound healing models. In this regard, it is noteworthy that several studies that have determined gentamycin to be less active than tobramycin [38, 40].

## Conclusions

In the present study we have demonstrated that tobramycin, although a less potent bactericidal compound *in vitro* than both moxifloxacin and ciprofloxacin, may be an effective countermeasure against *P. aeruginosa* infections through the deregulation of the T3SS secretion pathway and thus limiting the bacteria’s cytotoxic potential. The fluoroquinolones moxifloxacin and ciprofloxacin, on the other hand, appeared to potentiate T3SS mediated cytotoxicity at lower concentrations. These results could indicate that, when challenged by aminoglycosides, *P. aeruginosa* are less cytotoxic and invasive, with reduced capacity for systemic spread of infection. ExoU and ExoS expressing *P. aeruginosa* from bloodstream isolates of patients with bacteraemia were distinguished to be more susceptible to aminoglycosides amikacin (100% susceptible) and gentamycin (89% susceptible) than ciprofloxacin (48% susceptible) [12]. Aminoglycosides are sometimes administered to patients with another class of antimicrobial, such as a beta-lactam, in a combinational therapeutic approach [40]. Although we did not investigate beta-lactams on TS33 or antimicrobial combinations, the results of this study would suggest that a combination of a more bactericidal antimicrobial and a T3SS inhibiting aminoglycoside such as tobramycin, might serve to improve disease treatment outcome. In a study of combination antibiograms, to assess the susceptibility of *P. aeruginosa* from respiratory cultures, it was revealed that beta-lactam susceptibility ranged from 58% to 69% and addition of a fluoroquinolone or aminoglycoside resulted in improved in susceptibility. Importantly, however, only addition of tobramycin or amikacin provided susceptibility rates approaching or exceeding 90% [40]. Future studies should aim to assess whether or not tobramycin (and other aminoglycosides) can subvert the cytotoxic and proinflammatory effects of the T3SS in animal models of *P. aeruginosa* infection. Results of which could be pivotal in directing courses of medical treatment as well as giving us vital insights into other potential mechanisms by which antimicrobials can reduce the toxicity of bacteria.

## Acknowledgments

We thank Professor Dara Frank for kindly providing us with PA103 and the PA103 ΔUT mutant strain.

---

**Supplementary figure 1:**
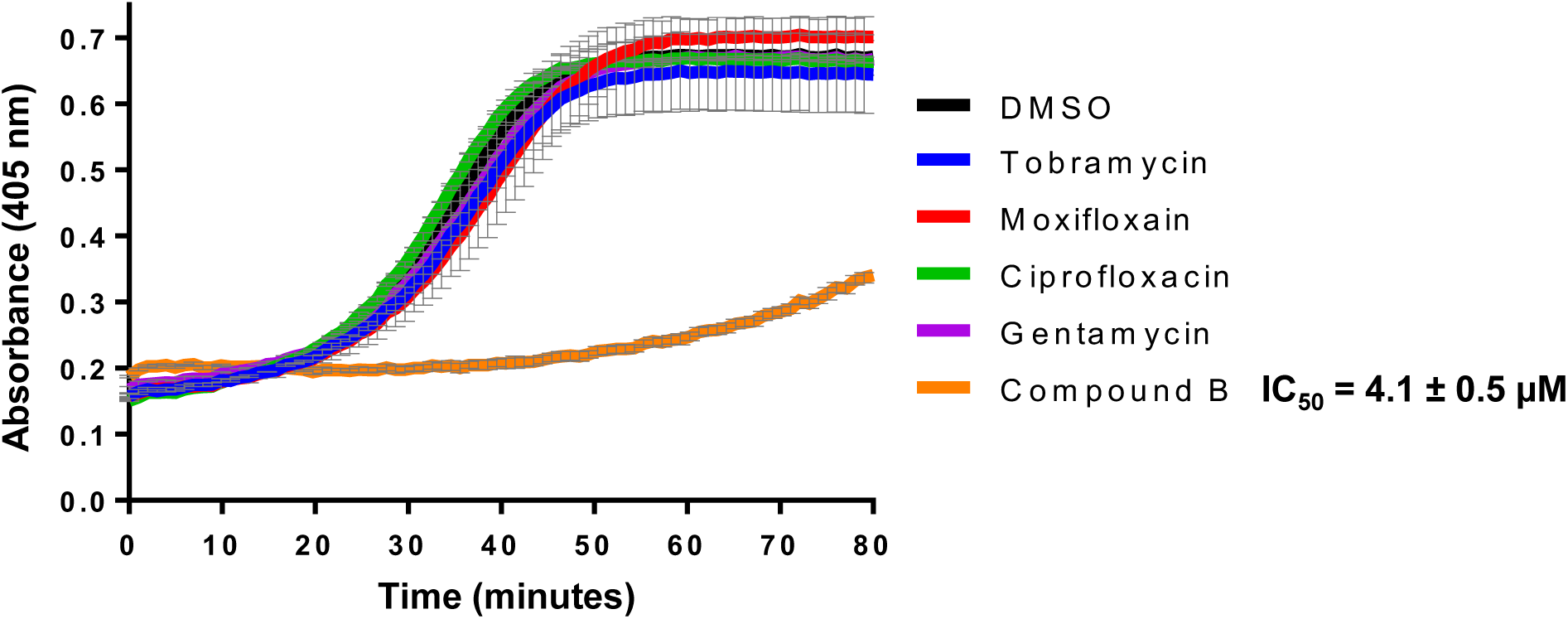
Antibiotics do not directly inhibit ExoU catalytic activity. The hydrolysis of arachidonoyl Thio-PC substrate by ExoU was assessed in the presence of 10 μM of antibiotic or compound B (positive control). To each reaction, ubiquitin and PIP_2_ were added for induction of ExoU phospholipase activity. Experiments were performed in triplicate, the results represent means, and error bars represent standard deviations and representative profiles are shown of substrate conversion as a function of absorbance with the progression of time.

**Supplementary figure 2:**
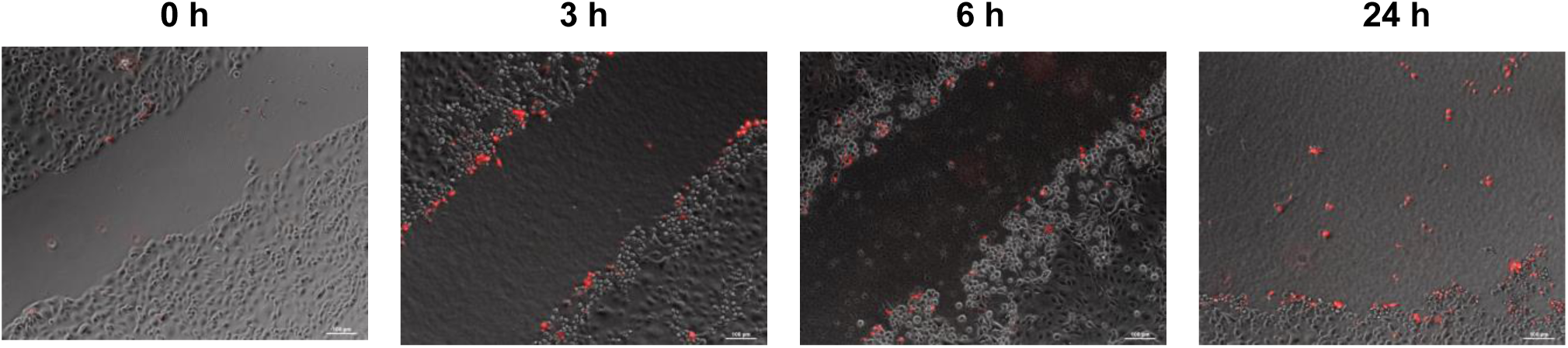
ExoS expressing PA76026 infection causes HCE-T cell rounding at the scratch boarder. Fully confluent HCE-T cells were scratched and then infected with PA76026 (MOI 2.5) for the indicated time points and analysed by microscopy. Cells were also stained with red fluorescent ethidium homodimer-1 to detect the presence of lysed cells.

**Supplementary figure 3:**
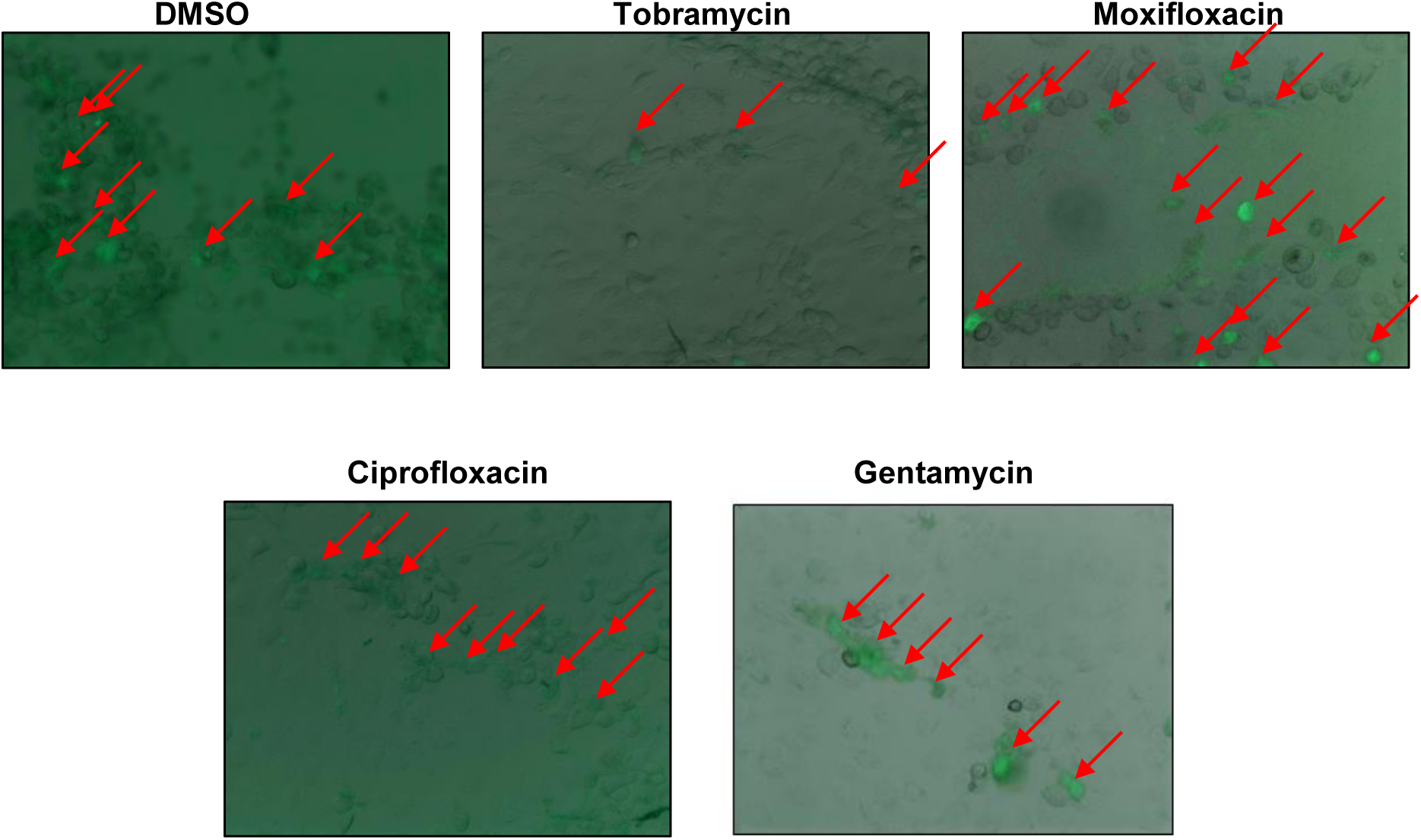
Intracellular GFP expressing PA76026 detected in HCE-T cells by fluorescence microscopy.

